# Natural slab photonic crystals in centric diatoms

**DOI:** 10.1101/838185

**Authors:** Johannes W. Goessling, William P. Wardley, Martin Lopez Garcia

**Affiliations:** Natural and Artificial Photonic Structures and Devices Group, International Iberian Nanotechnology Laboratory, Av. Mestre José Veiga, Braga 4715-330, Portugal

**Keywords:** Natural photonic crystal, Photonic band gap, Pseudogap, Biological photonics, Nanoporous silica, Diatom frustule, Diatom Girdle

## Abstract

Natural photonic crystals can serve in mating strategies or as aposematism for animals, but they also exist in some photosynthetic organisms, with potential implications for their light regulation. Some of the most abundant microalgae, named diatoms, evolved a silicate exoskeleton, the frustule, perforated with ordered pores resembling photonic crystals. Here we present the first combined experimental and theoretical characterization of the photonic properties of the diatom girdle, *i.e*. one of two structures assembling the frustule. We show that the girdle of the centric diatom *Coscinodiscus granii* is a well-defined slab photonic crystal, causing, under more natural conditions when immersed in water, a pseudogap for modes in the near infrared. The pseudogap disperses towards the visible spectral range when light incides at larger angles. The girdle crystal structure facilitates in-plane propagation for modes in the green spectral range. We demonstrate that the period of the unit cell is one of the most critical factors for causing these properties. The period is shown to be similar within individuals of a long-term cultivated inbred line and between 4 different *C. granii* cell culture strains. In contrast, the pore diameter had negligible effects upon the photonic properties. We hence propose that critical parameters defining the photonic response of the girdle are highly preserved. Other centric diatom species, i.e. *Thalasiosira pseudonana, C. radiatus* and *C. wailesii*, present similar unit cell morphologies with various periods in their girdles. We speculate that evolution has preserved the photonic crystal character of the centric girdle, indicating an important biological functionality for this clade of diatoms.

## Introduction

Photonic crystals (PhCs) - nanostructures with periodic features on the wavelength scale of light - have a wide range of applications in optical technologies, including high power lasers and quantum logic devices (1–3). Recent observations confirmed that PhCs evolved in nature, where they have been described for different biological phyla within the animal kingdom, foremost in invertebrates (4) and some vertebrates (5). Some recent discoveries demonstrated the presence of PhCs in photosynthetic organisms, including vascular plants (6) and macroalgae (7). Some of the most abundant microalgae, named diatoms, have attracted microscopists and scientists for centuries with the manifold optical phenomena occurring within their perforated exoskeletons, the frustules (8). This manuscript explores the PhC behavior of the frustule of centric diatoms, a significant clade of these photosynthetic organisms.

Diatoms are responsible for a quarter of global carbon fixation by photosynthesis (9). More than 100 000 different species, which can vary significantly in their shape and structure, populate almost all aquatic and most moist terrestrial environments (10). Two main morphological categories can be differentiated by their overall shape, determined by the frustule, i.e. a silicate exoskeleton surrounding the living cell. Centric diatoms are round, showing radial symmetry, while the pennate diatoms are elongated with lateral symmetrical frustules. The frustule is ornamented with micro-/nanometric pores, often highly ordered and reminiscent of the periodic cavities characteristic for some PhCs. Numerical analysis suggests that the porous network could show PhC properties in some species (11, 12); however, despite several studies that investigated the optical properties of frustules in different diatoms and with various techniques, to date, there has been no clear experimental evidence for PhC behavior of the frustule.

In both diatom categories, the frustule can be simplified as a construct made of two structurally different pieces: i) two convex shaped structures (valves) - each possessing slightly different diameters, making one fitting inside the other; ii) the valves are encircled by an even number of strip-like structures (girdles), located at the overlapping regions to keep the frustule exoskeleton densely stacked (Fig. 1 A). The frustule completely encloses the diatom cell, defining its shape and size (13). Because new valves form during asexual cell division inside the parental valves, the frustule slightly decreases in diameter following each asexual reproduction cycle. Once the frustule reaches a minimum cell size, the cell undergoes meiotic division involving a frustule-less stage, thereby reinstalling a maximum cell size of the species (14).

**Fig. 1:**
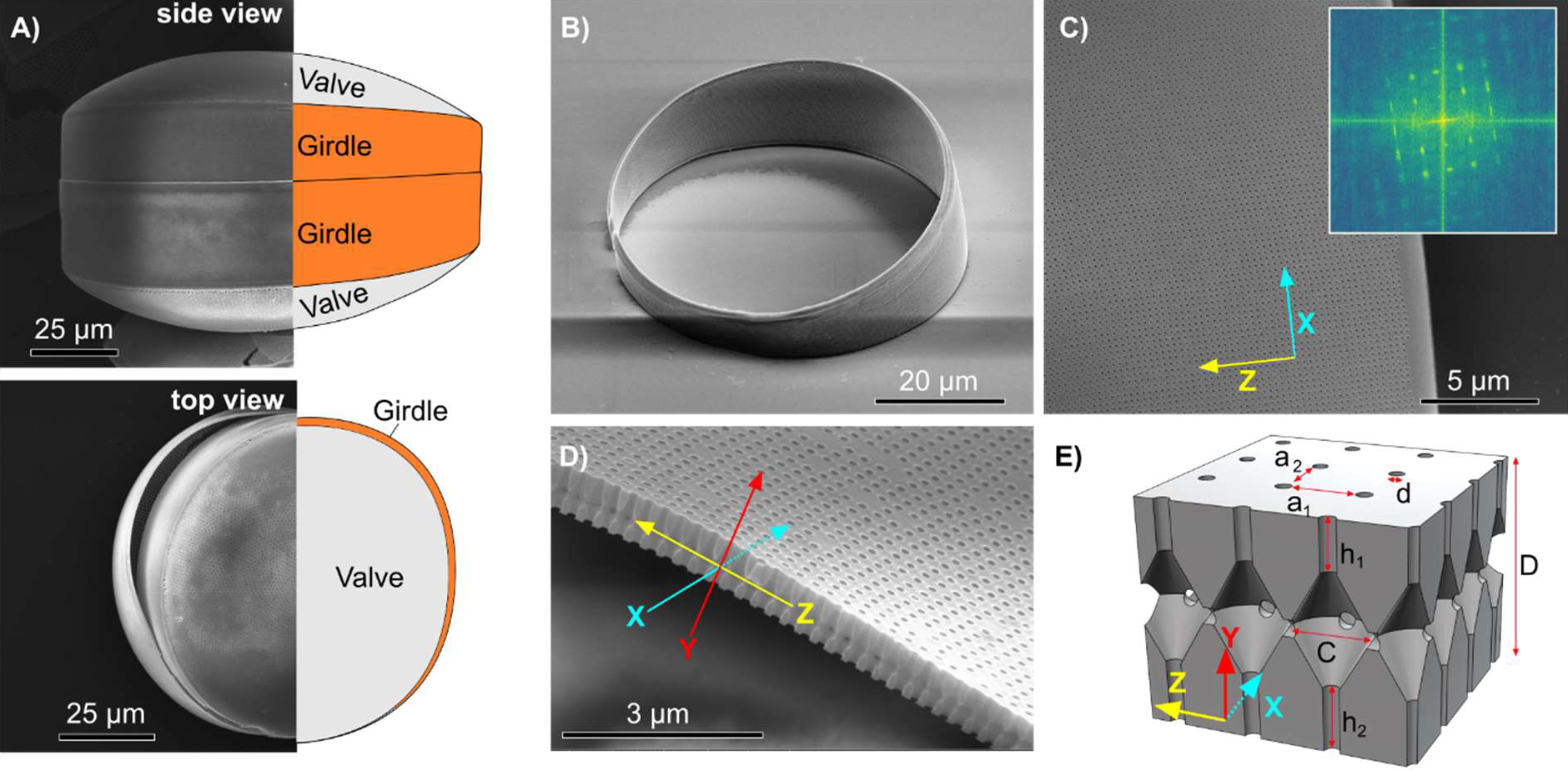
Frustule structure and girdle morphology of the diatom *C. granii* in SEM micrographs. The frustule of this species contains four silicate parts, i.e. two valves and two girdles. The girdles encircle the valves at the overlapping regions keeping the frustule together. A) Frustule from the side and from the top, with a girdle band visibly separating on the left side. B) Overview demonstrating the hollow cylinder character of the silicate girdle and the split ring spacing (on the left side). C) Surface micropores of the girdle in square lattice arrangement. Inset: Fast Fourier transform analysis of the lattice from the micrograph, demonstrating high periodicity of the pore arrangement. D) Cross section showing internal structure of the girdle along the Z axis. E) CAD reconstruction of the girdle crystal structure over multiple unit cells. Letters indicate the lattice parameter used in the optical model.

The silicate frustule is unique to diatoms and could have influenced their global abundance and species diversification (15). However, potential biological functionality of the frustule for the organism is hotly debated and remains elusive (16). One hypothesized function is the protection of the cell against micrograzers, as the frustule provides high mechanical strength combined with low Young’s modulus of elasticity (17). The frustule also facilitates cellular conversion of bicarbonate to CO_2_, promoting photosynthesis in aquatic environments by acting as a diffusion barrier (18). The pores allow for chemical communication with the environment, but may simultaneously prevent harmful agents like bacteria or viruses from entering the cell (19). The cause of the ordered arrangement, often in quasi-periodic, sometimes highly periodic arrangements, is unclear, but, as the period of structures and biologically useful sunlight wavelength scales matches, there is significant speculation that the frustule perforation ultimate function in modulating light-cell different interactions, e.g. by acting as a diffraction grating (20) or as a PhC (11).

PhCs are usually formed by a high refractive index material with a periodic structure at working wavelength scale (1). The patterning induces wave interference, building the optical response of the structure. The optical properties of PhCs are therefore strongly coupled to their morphologies (21). After the detailed investigation of PhCs over the last two decades, researchers have found a full zoo of structures (both natural and artificial) with a variety of exotic optical properties, not achievable with material properties alone. From a technical point of view, some of the most advanced PhCs are those referred to as slab PhCs, as they are complex multi-dimensional structures with specialized properties (22). Slab PhCs consist of micron-thick slabs of a dielectric material over which a periodic structure with different refractive index is induced. As the slab acts as a waveguide, the periodic nano-structuration can modify light propagating, causing photonic band gaps, i.e. directions for which light of a given wavelength cannot propagate. Slab PhCs are increasingly common in cutting edge technologies, ranging from sensing (23) and low footprint lasers (24) to quantum technologies (25). The photonic systems found in nature that have been studied to date, typically feature one dimensional multilayer structures (26) or complex 3 dimensional (3D) photonic structures (27); demonstration of natural slab PhCs (also known as 2.5D PhCs) has so far been elusive.

Previous studies into the photonic behavior of diatoms have investigated the optical and potentially photonic character of valves in different diatom species. Experimental studies on centric diatom valves suggested higher transmittance of longer wavelengths in the red spectral range, matching the absorbance maximum of chlorophyll-a (28). Another study found high absorbance of blue light in a centric diatom frustule (29). It was also suggested that the frustule attenuates ultraviolet (UV) light for photoprotection (30). Others found enhanced forward scattering of blue light in the valves of the centric diatom species *Coscinodiscus granii*, and speculated that these wavelengths can provide more energetic radiation for supporting photosynthetic electron transport. The latter study also found experimental evidence for photonic waveguiding within the valve (31). The valve of this species is formed by pores of different sizes and honeycomb chambers in hexagonal arrangement, with a period of ≈ 1250 nm and relatively non-uniform features. This suggests that interaction with visible light is driven by local diffraction, rather than PhC properties. In fact, the valve of different centric diatoms species seem to function as a diffraction grating, potentially enhancing or reducing light by interference at distances where the chloroplasts are located (32).

Regardless of its potential morphological importance, the girdle is a largely understudied part of the frustule in terms of its optical properties, as almost all experiments concerned the diffractive effects of the valve. In the few studies observing the girdle optical properties, this frustule piece has been described as a structure formed by a square lattice of cylindrical holes perforating a silicate slab (11, 31). The photonic properties of the girdle were only theoretically investigated and described as a slab PhC (11), where the dispersion relation of the guided modes is tailored by the photonic environment to form photonic bands (see 22). To date there was no clear experimental evidence for PhC properties of the diatom frustule. Consequently, potential biological functionality of PhC properties in the diatom frustule remained purely speculative. The lack of experimental evidence for PhC function also hampered the exploitation of diatom frustules for bio-inspired technologies. Combining a full description of its inner morphology with structural SEM analysis, Fourier micro-spectroscopy for optical characterization and theoretical models, the current manuscript demonstrates that the centric diatom species *Coscinodiscus granii* evolved well-defined slab PhC structures in their girdles. We also show that similar structures exist in the girdles of other centric diatom species, inferring evolutionary preservation and thus, potentially important biological functionality of the girdle PhC.

## Results and discussion

As well as the small number of studies on the optical properties, earlier morphological characterization of the girdle have also lacked important details (11, 33), i.e. the inner 3D morphology and material properties of the silica slab including nanoporosity. As we will expand upon through this manuscript, such details are essential for building the unique photonic response of the *C. granii* girdle.

Fig. 1 presents the overall shape of the centric diatom girdle as a circular silica slab, slant cut towards a split ring spacing (Fig. 1B). The radius of the slab depends on the cell diameter, which can vary in the approximate range 40 to 200 μm (34). The height of the slab within one individual girdle differs, as it tapers towards the split ring space. The thickness of the silica slab measured from SEM micrographs was D ≈ 745.1±42.7 nm (Tab. 1). The photonic crystal character of the girdle has been earlier attributed to a square lattice of micropores, perforating the silica slab in Y-direction with defined pore diameter *d* and lattice parameter a (11, 33). Fast-Fourier-Transform analysis over SEM micrographs (Fig. 1C) confirmed the well-defined square lattice of micropores, with similar period (P=0.257; N=10) along the X- and Z-direction, i.e. a_1_ = 284.7±4.8 nm and a_2_ = 279±11.0 nm, respectively. But analysis over the 3D inner morphology revealed strong differences compared to assumptions communicated for this species before (11, 33). In fact, a set of cylindrical micropores in X- and Z-directions intersect a central rhombic chamber, with the micropores inter-connected along the entire girdle slab (Fig. 1D/E). Based on the structural dimensions of the unit cell defined in Table 1, we calculated that the total volume occupied by micropores and rhombic chamber can account for ≈ 25-30% (see methods) in the unit cell. This volume defines the void filled with the surrounding medium, causing refractive index contrast.

In addition to the refractive index contrast between micropores and silica, the material properties of the silica slab will also affect the photonic characteristics. Earlier studies assumed a constant refractive index of the silica slab, usually n_silica_ = 1.45 (11, 12). This is possible, when amorphous silica is assumed for the diatom frustule (35). However, it has been communicated that the frustule could in fact bear nanoporosity (36). The girdle could therefore inherit a combination of two types of pores: the micropores that form the periodic structure, in addition to nanopores that characterize the material properties of the silica slab. Fig. 2A shows a SEM micrograph where such nanoporosity can be appreciated, followed by a sketch of a girdle unit cell, illustrating these two different pore types (Fig. 2B). The aspect is important for the optical description, because nanoporosity influences the effective refractive index of the silica slab, as the surrounding medium could penetrate into the nanopores (37). We address this important aspect during the following paragraphs more in detail, when measurements and modeling of the girdle photonic properties are presented.

**Fig. 2:**
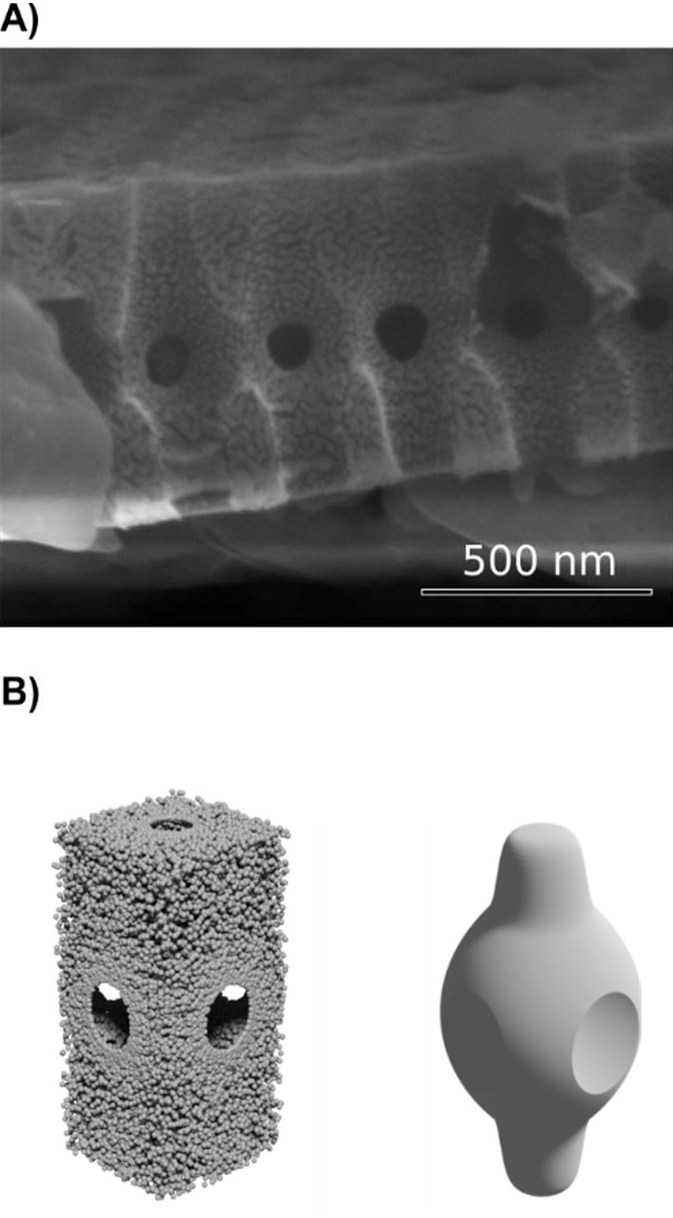
Micro- and nanoporosity defining the void filling volume in the *C. granii* girdle. A) SEM micrograph of a girdle cut along Z-axis. The micrograph indicates nanoporous characteristics of the bulk silicate skeleton. B) CAD illustration of the unit cell including nanoporosity and of the void presented by micropores.

**Table 1:**
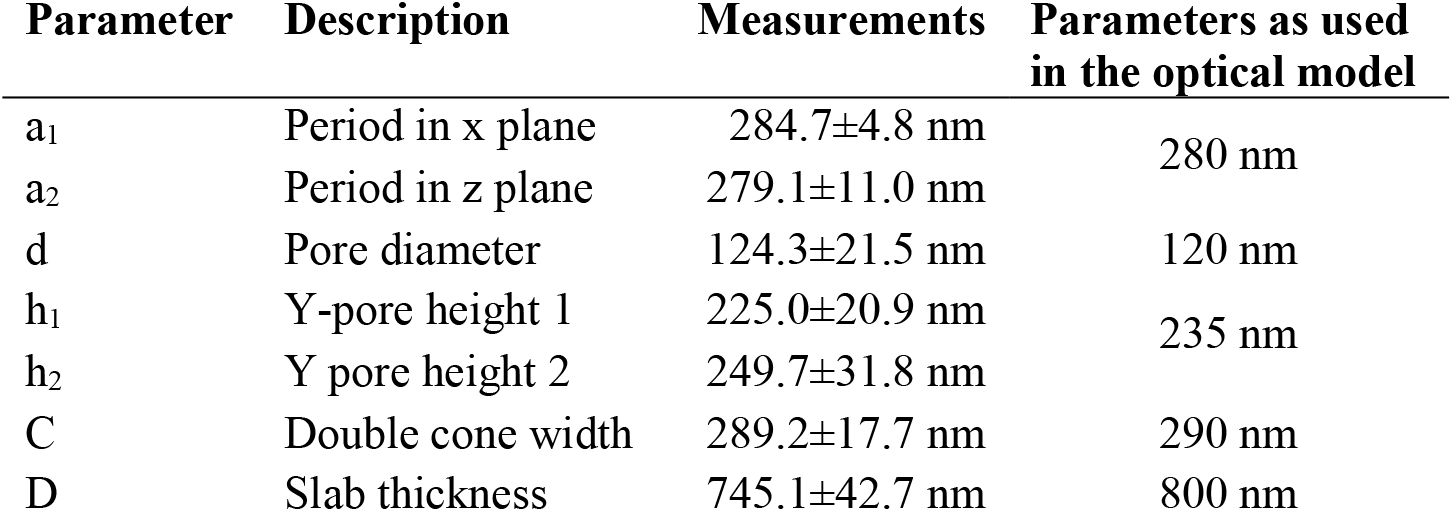
Unit cell characterization of the *C. granii* girdle. Measurements were performed on SEM micrographs, representing measurements of 5 individual girdles of the diatom *C. granii* (strain K-1834). Periods (a_1_ and a_2_), as well as pore diameter (d), were determined on 10 individual SEM images of the same diatom strain. Parameters used in the optical models are indicated.

| Parameter | Description | Measurements | Parameters as used in the optical model |
| --- | --- | --- | --- |
| $a_1$ | Period in x plane | $284.7 \pm 4.8$ nm | 280 nm |
| $a_2$ | Period in z plane | $279.1 \pm 11.0$ nm | |
| $d$ | Pore diameter | $124.3 \pm 21.5$ nm | 120 nm |
| $h_1$ | Y-pore height 1 | $225.0 \pm 20.9$ nm | 235 nm |
| $h_2$ | Y pore height 2 | $249.7 \pm 31.8$ nm | |
| $C$ | Double cone width | $289.2 \pm 17.7$ nm | 290 nm |
| $D$ | Slab thickness | $745.1 \pm 42.7$ nm | 800 nm |

To probe for photonic properties of the girdle, we measured reflectance of a specimen lying along the Z-direction normal to the glass cover slide (Fig. 3A). By this, we ensured coupling to the confined guided modes along Z-direction, as this would be expected for slab PhCs (38). Focused white light illumination, incident from Z-direction, would then correspond to in-plane waveguiding, facilitating high symmetry directions of the PhC (Fig. S1). Fig. 3B shows the well-defined reflectance on a girdle immersed in water, peaking at the central wavelength (λ_c_) ≈ 780 nm (near infrared; NIR) at normal light incidence (see also Fig. S2 with total reflectance values). The high reflectance and well-defined shape of the central peak, confirmed high degree of lattice order at optical wavelength scales. We then fitted the experimental measurements with Finite Difference Time Domain (FDTD) analysis. This approximation allows prediction of reflectance, when the introduced parameters match the effective indices of bulk material, void filling and surrounding medium, and material properties such as nanoporosity as well as the morphological characteristics. To obtain high agreement between measurements and model, we introduced the morphological parameters from Table 1, the refractive indices of water (n_water_ = 1.33) and bulk silica (n_silica_ = 1.45), and considered δ_i_ = 0.05, with δ_i_ describing the void volume introduced to the silica slab by nanoporosity. By this, the effective refractive index of the silica slab was decreased to n_silica_eff_ = 1.44 in water and n_silica_eff_ = 1.43 in air. We concluded that the strong central reflectance in the NIR indicated a pseudogap of the photonic bands, i.e. a photonic band gap that only exists for propagation in particular directions within the PhC structure (39).

**Fig. 3:**
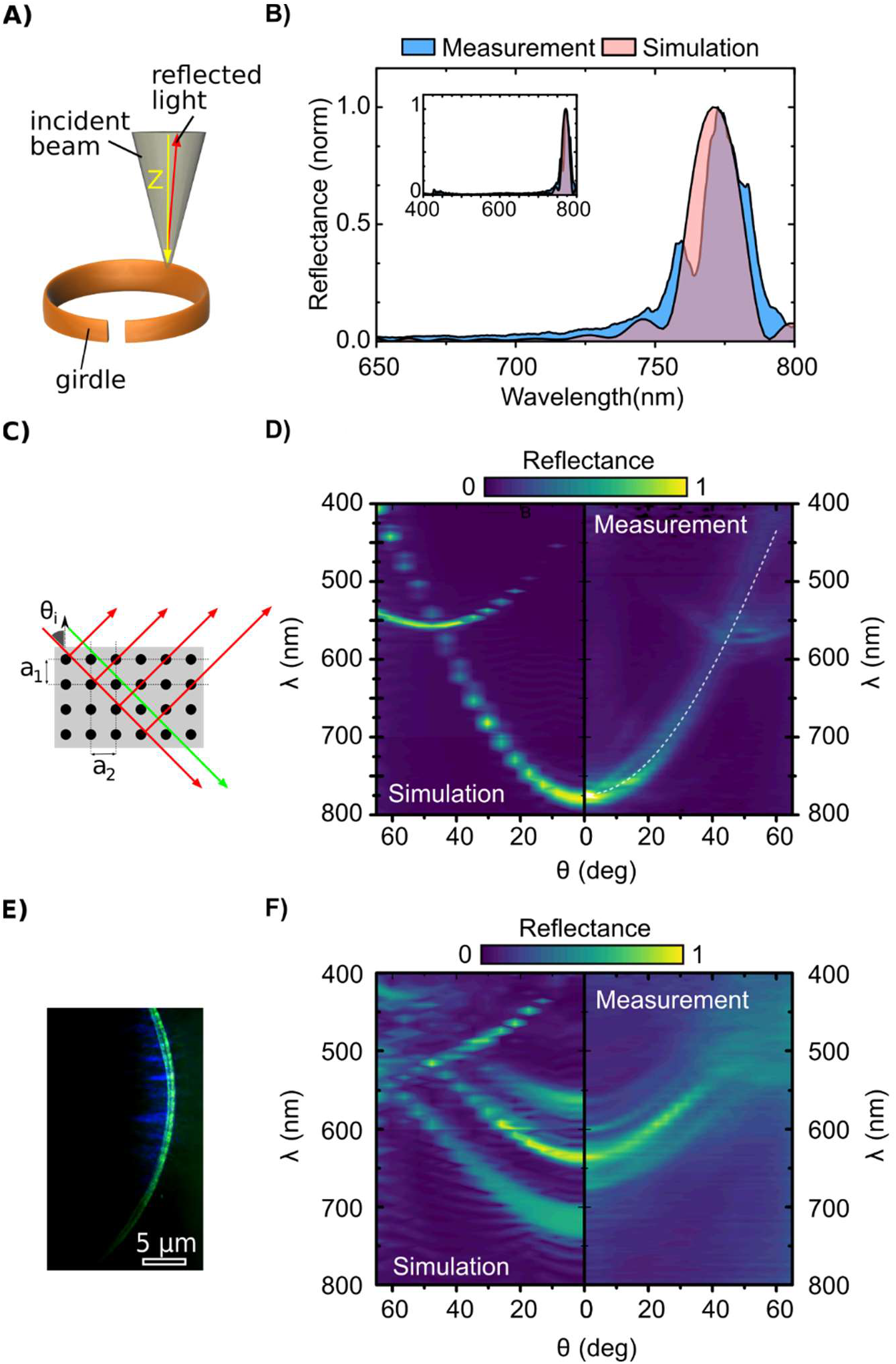
Photonic properties of the *C. granii* girdle. A) Sketch of the experimental setup, describing the direction of the focused white light (≈2μm) on the girdle (orange) in Z-direction. B) Experimental and FDTD simulations for the reflectance at normal light incidence. The inlet shows reflectance over the entire measured wavelength range. C) Sketch illustrating how light interferes with the micropores in Z-direction. D) Reflectance as a function of angle of light incidence and wavelength, observed with an oil-immersed objective (100X) with large numerical aperture (NA=1.45). FDTD simulation (left) and measurements (right) are shown. The dashed white line represents the central wavelength of reflectance when the effective index approximation is applied using the parameters from Table 1. E) Green light reflection by in-plane guided modes as suggested earlier (6). Blue light is also visible as it diffracts on the pore lattice. F) Fourier-micro-spectroscopy measurements in air, including FDTD simulation (left) and measurement (right).

To investigate the pseudogap further, we measured the reflectance as a function of light wavelength and angle of incidence (θ_i_), and fitted the measurements with the effective refractive index approximation (40). Using this approach, the 2D lattice of pores in Y-direction can be described as a Bragg stack with period a_1_ (Fig 3C). The contour plot in Fig. 3D illustrates that the pseudogap blue shifted as θ_i_ increased. The dashed white line shows the conformity of the effective refractive index approach confirming the reliability of parameters introduced in the FDTD. In addition to the reflectance caused by the pseudogap, a secondary reflectance pattern appeared at λ ≈ 500 for θ_i_ ≈ 30^0^. This secondary reflectance pattern shifted towards the red spectral range and increased in its reflectance intensity towards a maximum at λ≈ 560 nm for θ_i_ ≈ 50^0^, where it crossed the pseudogap. The secondary reflectance pattern could be explained by in-plane diffraction of the guided modes over period a_2_ in X-direction, which is indicative of the 2D symmetry of the slab PhC lattice as well as a proof of the high quality of the natural lattice inspected here. To provide visual proof of this property, we observed a water immersed girdle, illuminated with large angles, while collecting reflected light with large numerical aperture lens (Fig. 3E). Results showed that the girdle band produced strong structural colors through in-plane diffraction in the visible spectral range under this particular illumination conditions. Similar microscopic observations were communicated, but could not be explained, in the few earlier studies concerning the *C. granii* girdle (11, 33). The validity of the theoretical approximations was then further tested on girdles in air (n_air_ = 1.00), in the same experimental configuration. As shown in Fig. 3F, blue-shifted optical features with a strong optical peak around λ_c_ ≈ 630 nm were observed. When compared with the simulation results, we observed that a second and third strong reflectance peaks was present whilst the longer wavelength peak is missing in the measurement (λ_c_ ≈ 740 nm). The latter can be attributed to the fundamental reflectance peak of the PhC whilst second and third peak (λ_c_ ≈ 630 nm and λ_c_ ≈ 570 nm, respectively) are induced by the strong refractive index contrast between air and silica, as compared to the case of water. Probably, the strong refractive index contrast is also responsible for the lack of signal to unveil the longer wavelength peak in the measured signal.

Although observation of such precise slab PhC in a natural system is astonishing, whether this diatom species requires photonic properties to perform in its environment remains open. It is, however, also interesting that the observed properties can occur under more natural conditions, when the girdle was submerged in water and the refractive index contrast within the slab PhC was therefore low. The *C. granii* girdle structure can be regarded as a phenological trait, and, in evolutionary theory, such a trait is preserved when it fits for the functional role it performs. To investigate the degree of preservation of the PhC of the *C. granii* girdle, we validated the optical measurements on a second *C. granii* strain. In Fig. 4A we show the high similarity of the angular dispersion of the pseudogap between the strains K-1834 and K-1831. We have obtained the same angular and spectral dependency for at least five girdle measurements, to conclude that the girdle photonic behavior is highly preserved between individuals and between the two *C. granii* cell culture strains. A further comparison, concluded in Fig. 4B, confirmed the high level of preservation for the lattice period (a_1_ and a_2_) between four different cell culture strains of *C. granii* (a_1_: P=0.244, a_2_: P=0.534; N=10); while simultaneously, the surface pore diameter *d* (Fig. 4C) could significantly differ between some cell culture strains (P=0.008; N=10). Using our optical model (effective refractive index approximation), we found that the pseudogap dispersion is strongly affected by variation in period a_1_, but nearly unaffected by variation of the pore diameter *d*(Fig. 4D). We concluded that period a_1_, as a critical parameter of the PhC character for causing the pseudogap, is highly preserved and speculate that such preservation could be indicative for biological functionality of the girdle slab PhC. We then also investigated the interspecies preservation of the slab PhC morphology, by observing the girdles of three further centric diatom species using SEM techniques. All species in our experiments showed surface pores in a similar way to *C. granii* girdles. Cross-sections furthermore confirmed similar inner morphologies, including a central chamber at the core of the unit cell (Fig. 5). Such internal structures were previously presented in micrographs of the species *C. wailesii* and *Thalasiosira sp*. without further morphological characterization (41, 42). Although experimental confirmation for these species girdles is pending, the high structural resemblance suggests likely comparable PhC properties as demonstrated for the *C. granii* girdles. We found, however, that the period of the unit cell varied in the range 281±8 nm, 235±11 nm and 332±25 nm between these species, i.e. *Thalasiosira pseudonana, C. radiatus* and *C. wailesii*, respectively (determined on single SEM micrographs). As the period tailors the photonic behavior of the girdle PhC, different photonic characteristics are as well expected between these species. Such variation of a phenological trait might be expected in response to different environmental conditions, leading to niche differentiation and species diversification (43).

**Figure 4:**
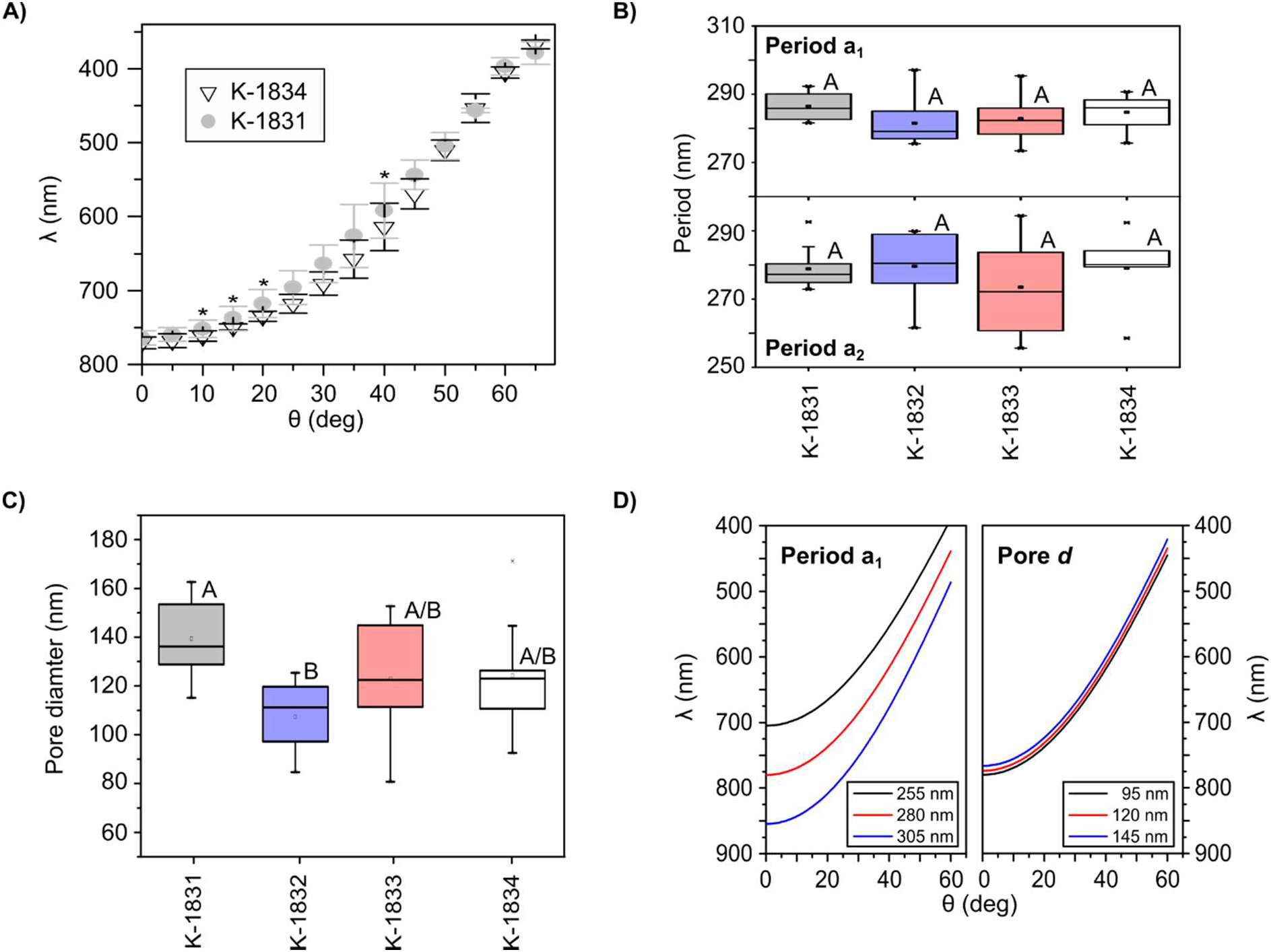
Comparison of PhC properties in different *C. granii* cell culture strains. A) Comparison of dispersion of the pseudogap in two strains of *C. granii* (K-1831 and K-1834) immersed in water. Significant differences, as determined with T-Test, are indicated with * at the P≤0.05 level. B) Comparison of the girdle lattice period (a_1,2_) in four cell culture strains of *C. granii*. C) Comparison of the surface micropore diameter in four strains of *C. granii*. Different capital letters in panel B) and C) indicate significant difference at the P≤0.05 level, determined with one factorial ANOVA and Holm Sidak posthoc tests. D) Effective refractive index approximation varying the period of the unit cell, or the surface micropore diameter *d*, respectively. Dimensional variation is indicated in nm.

**Fig. 5:**
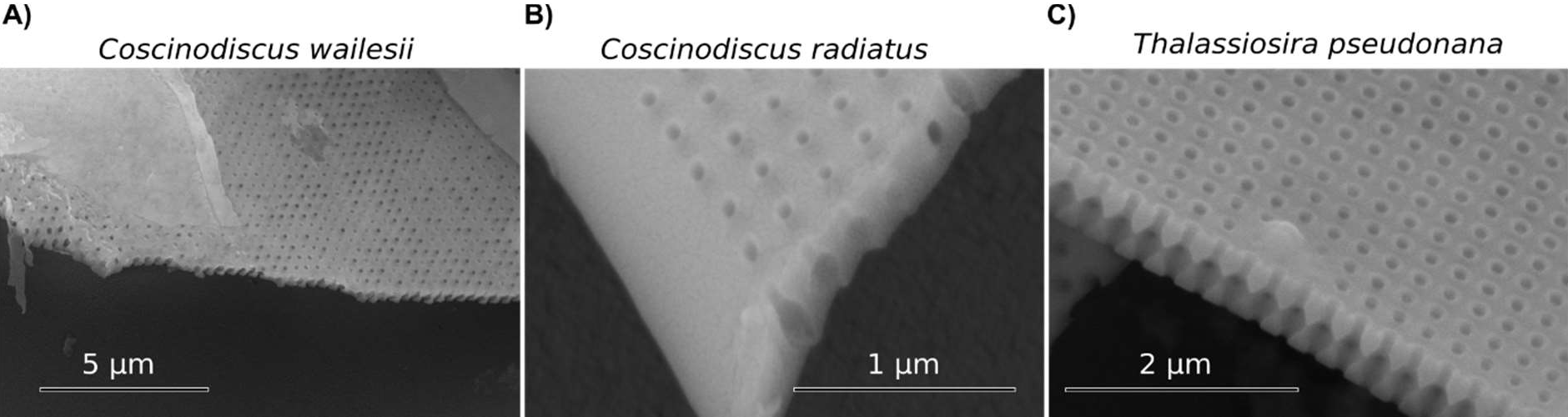
Internal girdle morphologies of different centric diatom species. All centric species tested in this study showed similar internal girdle crystallography, but varied in the period of the unit cell. A) *C. wailesii*, 332 ± 25 nm. B) *C. radiatus*, 235 ± 11 nm. C) *T. pseudonana*, 281 ± 8 nm.

In conclusion, we show that the girdles, parts of the *C. granii* frustule, are slab PhCs; to the best of our knowledge, this is so far the only slab PhC described experimentally for a natural system. It has been suggested that some plants and macro-algae use photonic structures to support their light harvesting and photosynthesis (6, 44). Earlier studies also discussed the possible role of the frustule related to its light modulating properties, mainly by facilitated passage for photosynthetically more productive (e.g. 28, 31, 45), or attenuation of potentially harmful radiation (e.g. 29, 46, 47). We here demonstrate the presence of a pseudogap with central reflectance in the NIR and waveguiding properties of green wavelengths under more natural conditions in water. Thereby, the PhC structure does not appear to be tuned for manipulation of photosynthetically more productive wavelengths in the species *C. granii*, as pseudogap and guided modes resonate in the range where light absorption for photosynthesis in diatoms is lowest (48). We speculate that the photonic response of the girdle could be involved in processes downstream photosynthetic light absorption, e.g. during energy dissipation or cellular capture of heat, or involved in perception of light. The girdle is an example of a very precise photonic material found in nature and can qualitatively compete with nanofabricated, artificial slab PhCs, opening the road for photonic-chip-like applications using naturally nanostructures. The apparent diversity of PhC properties (indicated by different periodicity of the PhC) within the centric diatom clade, may be further explored and could in future serve as inspiration for photonic innovation. Although some photonic systems in photosynthetic organisms have been described in the past few years, direct proof of their effects on photosynthesis, or on other physiological processes, remained elusive. However, the well-defined PhC property, the high structural preservation between individuals and different cell culture strains and the similar construction in other species, point towards significant biological functionality of the girdle slab PhC in the clade of centric diatoms.

## Materials and Methods

### Cultivation of diatom strains

Culture strains of *C. granii* (K-1831, K-1832, K-1833 and K-1834) and *Thalasiosira pseudonana* (K-1282) were purchased from the Norwegian Culture Collection of Algae (NORCCA). The species *Coscinodiscus wailesii* (CCAP 1013/9) was purchased from the Scottish Culture Collection of Algae and Protozoa (CCAP). Note that the diatom strains used in this study each originate from one single isolated cell, whereupon they proliferated by asexual and sexual reproduction during long-term cultivation in a culture collection. Diatom cell cultures were grown in L1 diatom medium (49) with a seawater base (30 ‰ salinity) and kept at a constant 18° C under low white light illumination (ca. 30 μmol m^-2^ s^-1^).

### Frustule preparation and removal of organic matter

Variable volumes (V) of diatom cultures were transferred to tubes. Initially, CaCO_3_ deposits were removed with 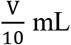 of 10% HCl. Afterwards, 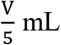 of 30% H_2_SO_4_ and 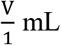 of saturated potassium permanganate were added and left overnight. Then, saturated oxalic acid was added until the mixture turned transparent. The mixture was centrifuged, before the supernatant was discarded and replaced with MilliQ water. This washing step was repeated thrice. Cleaned frustules were kept in MilliQ water until investigation. The cleaning procedure was adapted from (50).

### Electron microscopy and morphological analysis

Cleaned frustules were drop-cast onto silicon wafers and left to dry at 60° C, followed by 7 nm gold deposition with a multi-target confocal sputtering tool (Kenosistec, Binasco, ITA). The cover slip was mounted on a microscope stub and grounded with Electrodag silver paint. Frustule surface structures of 10 *C. granii* individual specimen of 4 strains (K-1831, K-1832, K-1833 and strain K-1834) were observed in non-tilted samples with a Quanta 650 FEG SEM (FEI, Oregon, USA), or with a dual beam focused ion beam SEM (FEI, Oregon, USA). The gold-covered frustules were shattered by pressing a glass cover slip sharply on top of the silicon wafer. By this, girdle fragments align normal to the stub with X- and Y-direction, facilitating observation of surface features in non-tilted mode. At the side of fracture, the internal morphology could be studied. Structures were analyzed on SEM micrographs using Fiji (51). Dimensions were aligned with a Pelcotec™ Critical Dimension Standard (AISthesis Products, Inc., Clyde, USA). Periods a_1_ and a_2_ and pore diameter *d* were measured over 20 surface micropores per girdle micrograph. In total, 10 individual girdles were measured for all *C. granii* strains, while one exemplary girdle was studied for each of the other centric diatom species. Measurements of internal structures were performed over 5 individual girdles for the species *C. granii*.

### Reflectance measurements with Fourier micro-scatterometry

Samples were characterized on an advanced Fourier image spectroscopy in a microscope setup adapted for the study of natural photonic systems (44). Frustule samples kept in water were drop-cast on a thin glass cover slip, and measured in water, or left to dry in an oven at 60°C for 24 h for measurements in air. A second glass cover slip was placed above of the sample using electrical tape as a spacer, to prevent large frustule pieces from breaking. White light illumination from a tungsten-halogen lamp was coupled with a 50 μm multimode optical fiber, then collimated and focused on the sample with a high numerical aperture oil-immersion lens (Nikon Plan-Apochromat 100×NA 1.45 oil OFN25). The beam was reduced to a spot diameter of ≈ 2 μm to probe reflectance from the girdle. Reflectance was collected for angles θ < asin(NA/n_oil_), with NA being the numerical aperture of the objective lens and n_oil_ the refractive index of the immersion medium. In our case NA=1.45 and n_oil_ = 1.51 therefore allowed for collection of θ_max_ = 74 deg. No movement or rotation of the sample was performed during measurement to ensure that the same volume of the sample was inspected for all angles. Reflected light was collected with a 100 μm optical fibre connected to a 2000+ Ocean Optics (Dunedin, USA) spectrometer. Each individual spectrum was normalized against the reflectance spectrum of a silver mirror measured under the same conditions. All spectral measurements were repeated on 5 specimens per group, if not stated otherwise.

### Simulation and modelling of photonic properties with FDTD

To model the photonic response of the girdle band we used commercial implementations of Finite Difference Time Domain Technique (Lumerical FDTD Ltd, Vancouver, CAN). The geometry of the internal girdle structure used for the simulations was obtained from the SEM analysis (Table 1). The selection of the refractive index parameters of the simulated structure considered a nanoporous nature of the material of the girdle, and is defined below.

### Approximation of nanoporosity

A table with abbreviations used for the refractive index approximations is provided in Table S1. The silica forming the frustule is assumed to be composed of SiO_2_ nanoparticles packed within a volume (17). The void between nanoparticles is filled with the surrounding medium of refractive index (n_i_). Therefore, the refractive index *n_silica_eff_* needs to be defined according to nanoporosity and fraction of volume occupied by the surrounding media of the biosilica (*δ_i_*). We use the Maxwell-Garnett approximation (52), which was validated for other natural nanoporous photonic structures (37). With this approximation, the effective dielectric constant of the biosilica 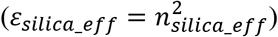 can be calculated, as:

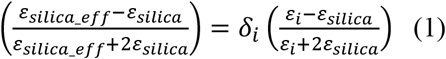

where 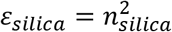 is the dielectric constant of bulk non-porous silica (n_silica_ = 1.45), ε_i_ = n_i_^2^ is the dielectric constant of the immersion material and *δ_i_* is the nanoporosity value (0 < *δ_i_* < 1). We consider n_i_ = 1.00 and 1.33 depending on whether the girdle is immersed in air or water, respectively. The nanoporosity value was determined by iteration of *δ_i_* in the FDTD reflectance calculation, until a maximum fit was achieved. By this, we obtained δ_i_ ≈ 0.05, which corresponds to effective refractive indices of n_silica_eff_ = 1.43 and 1.44 for a girdle band in air and water, respectively.

### Effective refractive index approximation for micropores

For the Bragg scattering approximation shown in Fig 3D and 3F, we used the effective refractive index approximation, validated previously for two dimensional hetero-structure slab photonic crystals (38). Each of the unit cells was considered as a scattering point arranged in the XZ plane (Fig. 3C). Applying Bragg’s law,

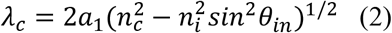

where λ_c_ is the central wavelength of the reflectance peak, a_1_ is the distance between planes which correspond to the modulus of the lattice vector, *θ_in_* is the incident angle respective to the normal plane of the photonic crystal and 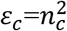 is the effective dielectric constants of the PhC, which has to be differentiated from the effective refractive index of the biosilica (n_silica_), as calculated before. This approximation relies on the effective refractive index of the PhC slab (n_c_) induced by the micropore void. Previous studies on complex 3D photonic crystals defined the effective refractive index of a photonic crystal, by taking into account the void filling fraction for a given material inside the unit cell (53). In the case of the girdle, the void filling fraction (f_i_) of the unit cell can be calculated as:

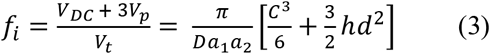

where V_DC_ and V_p_ are the volumes of the double cone chamber and X-,Y- and Z-pores, respectively. Note that we considered the interconnection of chambers by cylinders in X- and Z-direction with the same diameter, but, considering that C > h_1,2_ (Fig. 1D), the double cone chamber occupies the space of the X and Z-axis pores. All volumes are filled by the immersion medium (n_i_). The remaining volume is filled with silica (n_silica_eff_). V_t_ is the total volume of the unit cell with dimensions D and a_1,2_. Note that we considered a perfect square lattice (a_1,2_ = a), the X-, Y- and Z-pores with same dimensions, and the double cone chamber as a sphere for the sake of simplification. By this, we then defined the effective refractive index of the photonic crystal as:

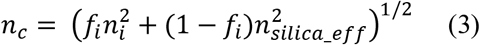

With this approximation, we obtain n_c_ = 1.25 and 1.39 for a girdle in air and water, respectively. Note that these values are dependent on ni, on which also n_silica_eff_ depends.

### Statistical analysis

Lattice vectors a_1,2_ in the *C. granii* strain K-1834 were tested with one-factorial analysis of variance (ANOVA) followed by Holm Sidak posthoc analysis. Comparison of four *C. granii* cell culture strains concerning differences in period a_1_, a_2_ and pore diameter *d* were treated in the same way. Reflectance data of K-1831 and K-1832 girdles were tested with one-tail, homoscedastic T-test at increments of 5° theta. Differences at the P>0.05 level are considered significant. Statistics were performed using the software Origin (OriginLab Corporation, Northampton, USA).

## Supporting information

2019_Goessling et al_Supplementary

## Acknowledgments

J.W.G acknowledges support and co-funding of the NanoTRAINforGrowth II program (project 2000032) by the European Commission through the Horizon 2020 Marie Sklodowska-Curie COFUND Programme (2015), and by the International Iberian Nanotechnology Laboratory. W.W and M.L.G would like to acknowledge the project POCI-01-0145-FEDER-031739 co-funded by Fundação para a Ciência e a Tecnologia and COMPETE2020.

