## Supplementary material for "Natural slab photonic crystals in centric diatoms": 2019_Goessling et al_Supplementary

**Supplementary information to**
**Natural slab photonic crystals in centric diatoms**

By Johannes W. Goessling, William P. Wardley, Martin Lopez Garcia

**Table S1: Abbreviations used for morphological lattice description and refractive index approximations**

| | |
|---|---|
| $\delta_i$ | Filling volume of the immersion media in the nanoporosity approximation |
| $f_i$ | Void filling fraction of the micropores |
| $n_i$ | Refractive index of the surrounding medium |
| $n_{air}$ | Refractive index of air (=1.00) |
| $n_{water}$ | Refractive index of water (=1.33) |
| $n_{silica}$ | Refractive index of the bulk silica material |
| $n_{silica\_eff}$ | Effective refractive index of the silica slab including nanoporosity ($\delta_i$) |
| $n_c$ | Effective refractive index of the PhC |
| $\varepsilon_{bulk}$ | Effective dielectric constant of the bulk silica material |
| $\varepsilon_{silica}$ | Effective dielectric constant of the silica slab considering nanoporosity |
| $\varepsilon_i$ | Effective dielectric constant of the surrounding medium |
| $\varepsilon_c$ | Effective dielectric constant of the PhC |

**Photonic bands of the girdle**

Although a full description in terms of photonic bands is out of the scopes of this work, we present as supplementary information the photonic bands calculation using the same simulation parameters described in Table 1 of the manuscript. We therefore consider $a_1$=285 and $a_2$=278 in this case with d= 100 nm. To calculate the photonic bands of the girdles in water, we used the plane wave expansion method implementation provided by the MPB software (22). Since we were mostly interested in the description of totally confined modes (the ones inspected in our experimental configuration) we can approximate the lattice as pure 2D system instead of a 2.5D photonic crystal. Therefore, the calculation of the photonic bands is performed considering silica material perforated by a 2D lattice of infinite rods filled by refractive index $n_i$ = 1.33. For the silica, we used the values provided by the Maxwell-Garnett approximation of the silica (see methods in the main text). Under this approximation $n_c$ = 1.44.

As can be observed, the photonic bands in Fig S2 shows the low energy pseudogap for $\lambda\approx$785nm, which is the same spectral range for which we report the experimental evidence in this work (see Fig 3).

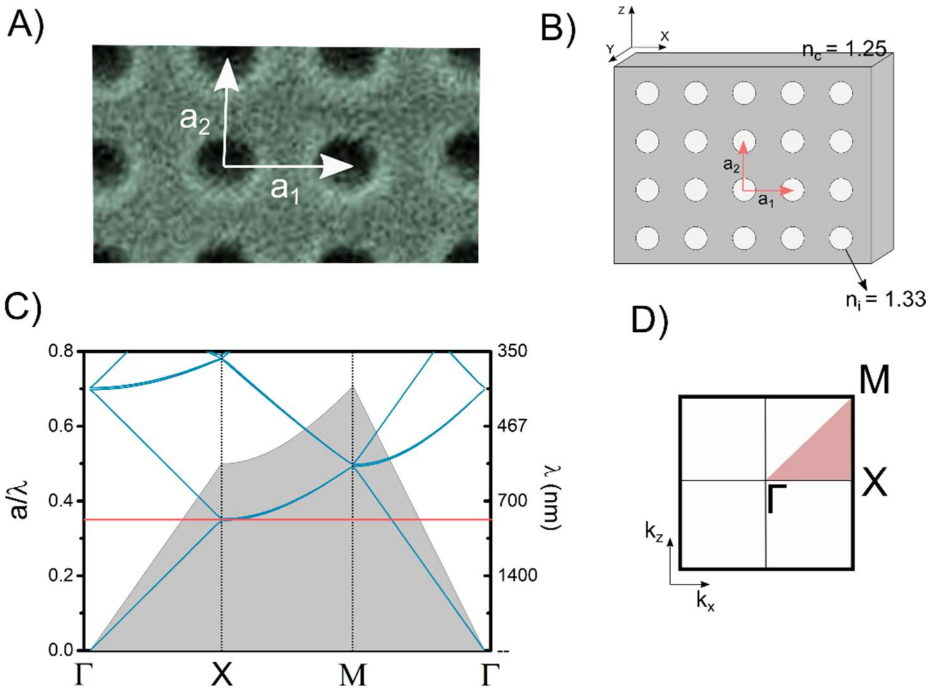


**Fig. S1:** A) Lattice vectors as defined over an SEM image of the girdle band photonic structure. B) Sketch of the lattice and refractive indices for bulk material ($n_c$ = 1.44) and rod filling material ($n_i$ = 1.33). C) Photonic bands calculation for TM modes. Red line indicates spectral position of the pseudogap. D) Brillouin zone for a square lattice.

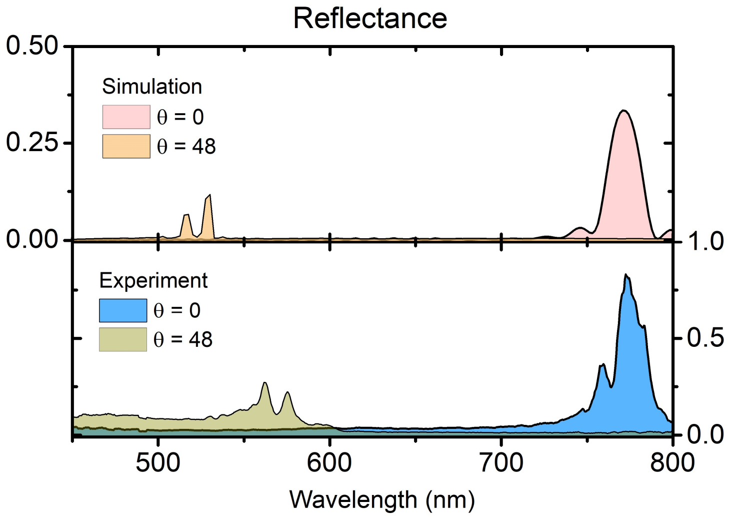


**Fig. S2:** Single absolute reflectance spectrum for two different incident angles and for the girdle in water shown in Fig 3D. Note that the values shown in the main text have been normalized to the maximum reflectance. The experimental absolute values shown here are the result of normalization to a silver mirror spectrum measured in the same conditions than the samples (see methods). The experimental values present larger values of reflectance than the predicted simulation, due to the complex illumination necessary to inspect the girdle in the appropriate lattice direction. However, the spectral shape of simulation and measurement are on a very good agreement both angles regardless of a slight detuning for θ = 48 deg.
